# Smart soils to observe hidden rhizosphere processes

**DOI:** 10.1101/2021.11.19.468202

**Authors:** Daniel Patko, Qizhi Yang, Yangminghao Liu, Panagiotis Falireas, Benoit Briou, Bhausaheb V Tawade, Timothy S George, Tim J Daniell, Michael P MacDonald, Vincent Ladmiral, Bruno Ameduri, Lionel X Dupuy

**Affiliations:** Ecological Sciences, The James Hutton Institute, Dundee, DD2 5DA, UK; Department of Conservation of Natural Resources, Neiker, Derio, 48160, Spain; Ingénierie et Architectures Macromoléculaires, Institut Charles Gerhardt, CNRS, University of Montpellier, ENSCM, Montpellier, 34000, France; School of Engineering, Physics and Mathematics, University of Dundee, Dundee, DD1 4HN, UK; Department of Animal and Plant Sciences, The University of Sheffield, Sheffield, S10 2TN, UK; Ikerbasque, Basque Foundation for Science, Bilbao, 48009, Spain

## Abstract

Agriculture must reduce green-house gas emission and pollution, produce safer and healthier food, closer to home, reducing waste whilst delivering more diverse diets to a growing world population. Soils could enable this transformation, but unfortunately, they have a hugely complex and opaque structure and studies of its myriad of mechanisms are difficult. Here, the fabrication of smart soils for the screening of below-ground bio-processes is demonstrated. Particles were generated from fluoropolymer waste with functionalisation converting them into sensors able to report on key chemical dynamics. Tailored functionalization was obtained by radical terpolymerisation to improve growth conditions and sensing capabilities. The study demonstrates the potential for the development of accelerated genetic or agrochemical screens and could pave the way for improved models for rhizosphere dynamics.

## Introduction

Crop production relies heavily on processes taking place within the complex microstructure of soils. Large variation in pore size buffer variation in rainfall and irrigation with small pores retaining water to maintain supply whilst large pores drain excess water^1^. Macro pores also promote gas exchange and respiration^2^, with these exploited by roots to avoid friction, waterlogging and excessive pressure from the soil^3^. The chemical complexity of soil particle surfaces favours adsorption of mineral ions, reducing leaching while slowly releasing nutrients for plant uptake^4^. Soils also provide the physical support for organic carbon deposition, transformation, and large-scale fixation^5^ and buffer, or filter, various forms of pollution^6^. The heterogeneity of soils is such that only a few grams of material hosts diverse microbial communities fulfilling a broad range of functions^7^, making it one of the most complex environments on the planet.

Agricultural production needs to increase by around 50% between 2012 and 2050 to achieve food security with degradation of soil quality recognised as a major constraint on this goal^8^. Soil is a non-renewable resource over a human time scale, but is being lost at a rate up to 100 faster than soil formation, with around a third of global soil reserves being lost in the last 50 years^9^. Significant sustainable gains in crop productivity can be achieved if we can successfully harness soil heterogeneity through a greater understanding of the regulation of its biological processes, making them more resilient to degradation. New technologies will be key but whilst tools for the proximal and remote sensing of soils are flourishing^10^ development for the monitoring of biological activity belowground remain limited. Remote sensing is often superficial because, for example, soil blocks electromagnetic signals in the visible domain and radio waves cannot resolve micrometric variation^11^. The application of penetrating radiations such as X-ray or neutron tomography successfully depict the physical structure of soils^12^, but cannot resolve functional biological or geochemical processes. Thus, the use of invasive approaches remains predominant in the soil sciences and because measurements permanently disrupt micro-habitats, our ability to understand, predict, manage and engineer soils for biological production remains limited.

Since *in vivo* measurements are rendered impractical in natural soil we asked the question, why not create soil-like materials that can report on the nature of the inner soil bio-processes? Recent studies have demonstrated the possibility of culturing and observing soil organisms in a transparent substrate that mimics natural soil properties^13–16^. The next logical step is to engineer the soil particles themselves to provide a more natural habitat enhancing growth whilst enabling the acquisition of various optical signals to sense soil conditions at a meaningful scale. Here we present the development of the first generation of such soils. We have optimised the smart soils for root growth and measurement of pH an important indicator of nutrient availability and plant stress. We demonstrate their fabrication from plastic waste products and assess how soil structure and root exudation contribute to the formation of chemical heterogeneity in soil.

## Results

### Functionalisation of Fluoropolymers allows fabrication of soil particles as sensors

Fluoropolymers have critical characteristics for application as optochemical sensing, e.g. they have both a low refractive index and high transparency^17,18^. They are also chemically inert and have properties such as photostability along with great chemical, thermal and oxidation resistance^19,20^. However some of these properties, such as hydrophobicity and chemical stability make them challenging to adapt as a soil substitute. Here, we explored using waste FEP (Fluorinated Ethylene Propylene, commonly produced by industries for tubing, coatings, and cable^21^) for the design of “smart” soil particles.

Waste FEP particles were functionalised with a thin polymer shell augmenting them with a range of custom properties (Figure 1). The design of the shell was achieved by conventional radical terpolymerisation of three comonomers, each of them bringing specific and complementary properties. We synthesised and trialled numerous materials (Supplementary Information 1) and identified key elements required for use as a plant growth medium. First, a fluoropolymer matrix was required for bonding, since it is known that Fluorine-Fluorine specific interactions are possible^22,23^. 2,2,2-trifluoroethyl α-fluoroacrylate (FATRIFE) monomers were used, but similar attachment could be obtained with hexafluoroisopropyl α-fluoroacrylate (FAHFIP). Plant growth also requires the particle to retain water and nutrients at the surface of the particle. Because of the hydrophobic behaviour of FEP and FATRIFE, such hydrophilicity was first achieved by addition of carboxylic acids 2-trifluoromethyl acrylic acid (MAF), methacrylic acid or Poly(ethylene oxide) groups^24^. Finally, various sensors can be included for acquisition of biological or environmental signals, and we demonstrate the development of sensing soils by focusing on pH sensing using Nile Blue (NBMA)^25^. The terpolymerisation of FATRIFE with polyethylene glycol methacrylate (PEGMA) and Nile blue methacrylate (NBMA), initiated by *tert-butyl* peroxypivalate (TBPPi), was achieved yielding a wide range of terpolymers of various compositions determined by ^1^H and ^19^F NMR spectroscopy.

**Figure 1.**
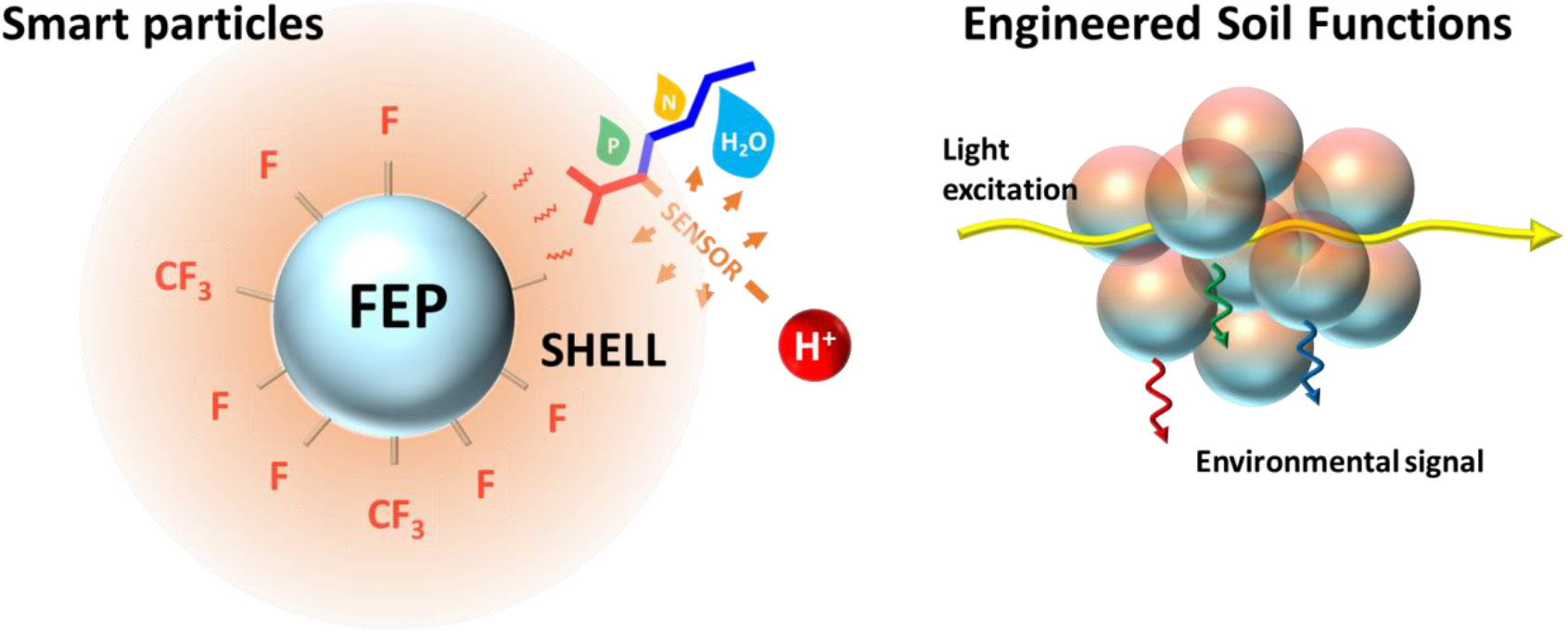
Engineering artificial functions for monitoring and controlling biological activity in soil. The soil particle is made of a core made of waste Fluorinated Ethylene Propylene (FEP), which is embedded into a functional material, the shell (left). The shell holds water and nutrients and includes a sensor (here a pH sensor). The soil is subsequently able to respond to external signals such as light of various wavelength and report on processes from the inner structure of the soil (right).

Because FEP is transparent and has a refractive index of 1.341 – 1.347^26^, the particle allows index matching in the soil interstices with water and subsequent acquisition of fluorescent signal from the inner soil structure. This process is similar to the use of optical clearing for imaging more deeply into human tissue using an index matching fluid such as glycerol. Here the particles themselves are engineered to allow for index matching with water rather than a higher refractive index fluid which would not support plant growth.

### Functional materials improve growth and allow attachment of pH sensor

In the following sections, we used the poly(FATRIFE-*ter*-PEGMA-*ter*-NBMA) terpolymer with 75/25/0.1 molar ratio (Figure 2A). The composition of the coating was characterised using Fourier-transform infrared spectroscopy (FTIR) which demonstrated successful deposition of the material on the surface of the particles (Figure 2B). Stability of the coating, which was tested on particles immersed in water and submitted to shaking for at least 4 weeks, demonstrated a strong adhesion on the FEP. The coating drastically improved the water interfacial tension of FEP particles and the ability of the soil to hold water. This was characterised using water retention curves (Figure 2C). The particles were subsequently tested for their ability to deliver optical signals using a light sheet microscope (Figure 2D). The soils were trialled for biocompatibility (Figure 3) and biomass production. Further details of polymer synthesis and the various methods used for characterisation are presented in (Supplementary Information 1).

**Figure 2.**
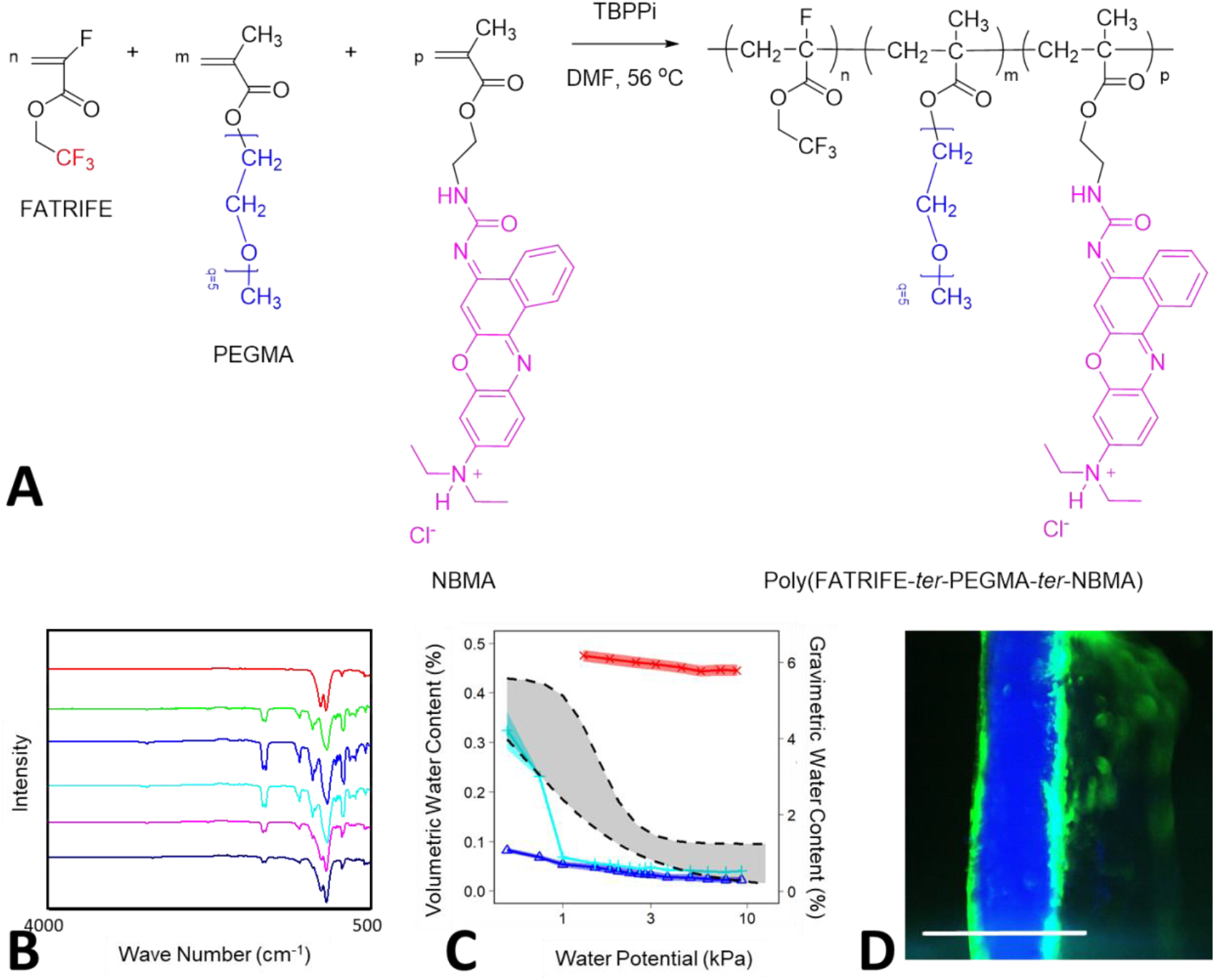
Using FEP particles for the fabrication of smart soil particles. (A) Chemical structure of the polymeric shell material encoding three different functions. The polymer consisted of Alpha fluoro-2,2,2-trifluoroacrylate (FATRIFE) for the attachment onto the FEP core, PEGMA enabled to enhance water retention and Nile Blue methacrylate (NBMA) brought the pH sensor. Materials were tested for various properties during the optimisation of the chemical structure (Supplementary Information 1). (B) The stability of coatings were studied using FTIR. FEP FTIR spectrum (top red) was compared to those of the coatings immediately, after 1 week, 2 weeks, one month and two months (from green top to bottom black respectively). (C) Results showed that terpolymers containing PEGMA (red x) increased the water retention of FEP soil particles (cyan +) with comparison to non-functionalised particles (blue Δ). Water retention curves were compared to those of sands (dashed line). Shaded areas indicate mean value ±SE for artificial soil and the range of values observed in natural sandy soils. (D) The particles were subsequently tested for imaging using light sheet scattering (blue) and fluorescence (green) signals (scale bar 1 mm).

**Figure 3.**
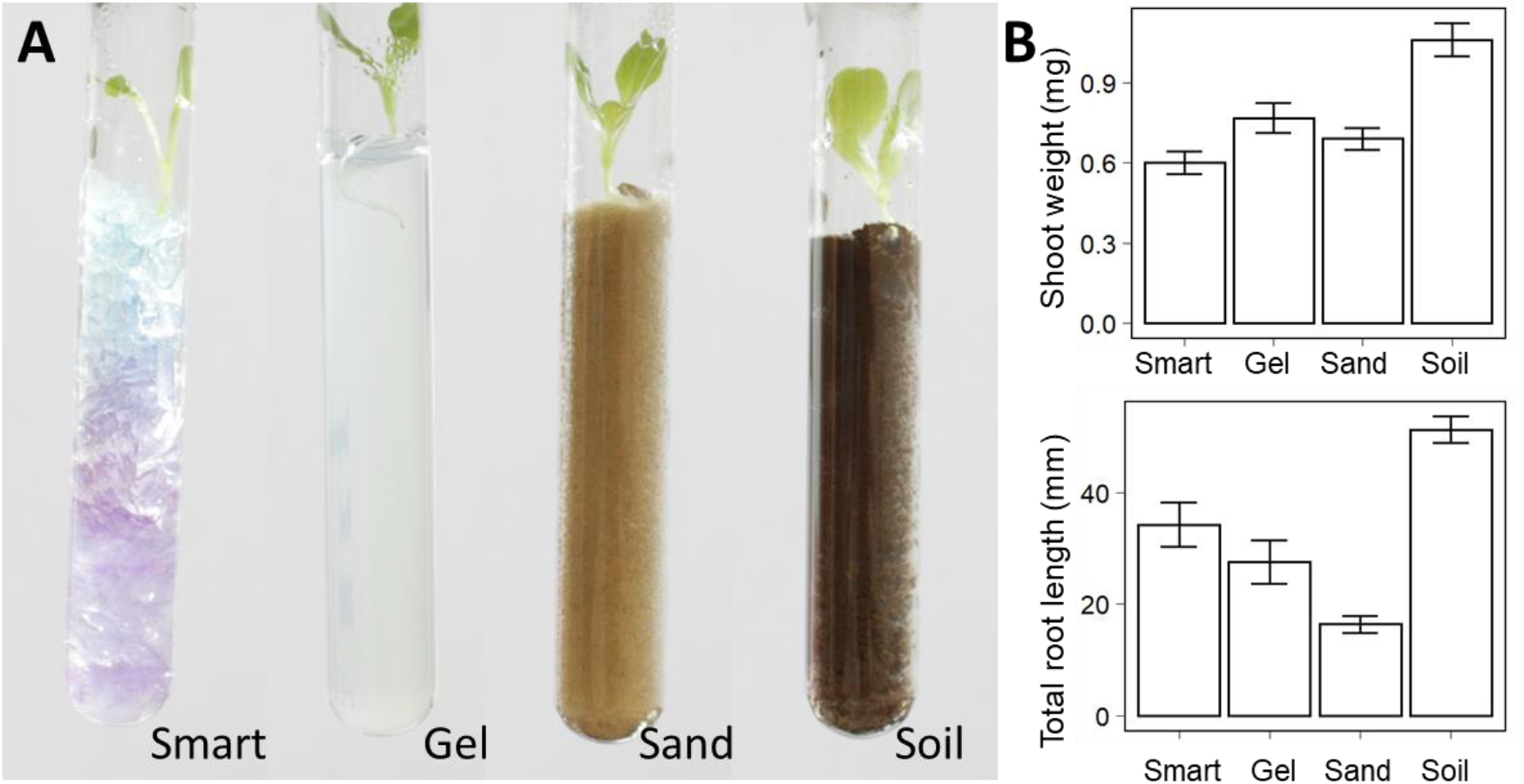
Biocompatibility of smart soil particles (A) Growth of lettuce seedling in smart soil particles, agar gel, sand and an agricultural soil. (B) Plants produced larger shoot and root biomass in an agricultural soil than in all other types of soils (p<0.001, n=10). Smart soil exhibited stronger root growth than other artificial media, although the difference was statistically significant only with comparison with sand (p= 0.001231, n=10). An analysis of variance showed both shoot weight and root length were significantly affected by soil treatments (p<0.001). Error bars indicate mean value ±SE for artificial soil and the range of values observed in natural sandy soils

### Multispectral light sheet imaging resolves pH changes at the micro-scale, live and in situ

We subsequently developed a multispectral light sheet imaging approach for measuring pH changes within soil samples. Although particles showed consistent changes in fluorescent intensity, the response varied significantly between individual particles (Figure 4A&B). To overcome this we first attempted a ratiometric fluorescence approach, but this yielded pH predictions with limited accuracy. The best results were obtained with light excitation at 488 nm and 633 nm and calculation of prediction intervals showed 95% predictions of pH values were within −0.6 to +0.5 of the true pH value. Machine learning proved more successful. The method used four illumination wavelengths (488 nm, 514 nm, 561 nm and 633 nm), for each wavelength a fluorescent signal was recorded and a neural network model was used to predict pH values from both the xy-position in the image and the collection of fluorescent responses from all four excitation wavelengths (Figure 4C). Results showed the machine learning approach improved predictions significantly in a test dataset (Figure 4D, t=79.9, p<0.001), with 95% of the predictions falling within −0.4 and 0.3 of the true pH value, almost doubling the precision of ratiometric fluorescence predictions. Ratiometric fluorescence predictions also introduced bias leading to underestimation of large pH values. Up to 80% of variations in pH were correctly predicted using neural networks and only 46% of the variance was explained by the ratiometric fluorescence approach.

**Figure 4.**
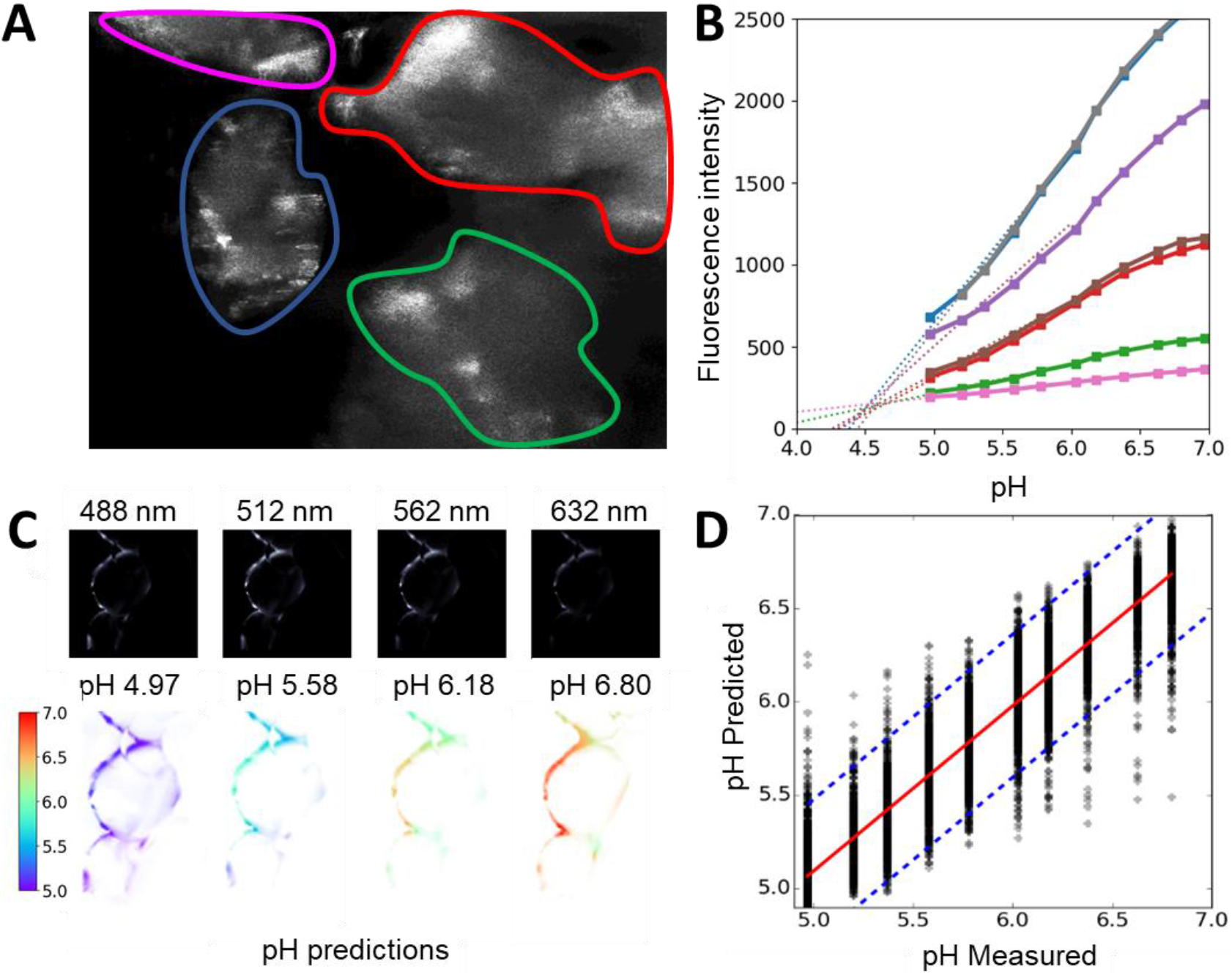
Calibration of pH sensing particles. (A) Smart soils imaged in a light sheet microscope contain scattering artefacts which reduce the power of ratiometric analyses. (B) Nevertheless, particles showed near linear increase in green fluorescence (488 nm / 530 nm) in response to pH between pH 5 and 7 (each colour represents a different particle in the same image). (C) A machine learning approach enabled the use of data from all fluorescent signals (top) to predict pH from image datasets (below). Pseudo colours indicate pH predictions corresponding to the pH buffer used (indicated above each image). (D) pH predictions using the machine learning approach explained up to 80% of the variance of the data and exhibited limited bias.

### Smart soil particles help understand the formation of soil micro-habitats

Using the fabricated soil particles, plants were observed up to one week after germination. The first sign of the pH change was observable after 4 days which is consistent with classical studies^27^. Following growth in the smart soils, we could observe the changes in soil chemical properties and determine how these changes relate to soil structure and root biological activity (Figure 5A). Maps of entire seedlings and surrounding soils were assembled by tessellating images to produce volume data corresponding to up to 3 cm^3^ of soil at 10 μm resolution with each time point producing a dataset of approximately 4 Gb at maximum dynamic range (16 bits image data) ^28^. pH predictions obtained from the data (Figure 5B-D) revealed how soil micro-habitats may form as a result of root exudation, and how this process is influenced by the generation of microsites and the type of FEP particles. Non-linear mixed effect models (Supplementary Information 2) showed that when roots grew in soil made of chunk FEP particles (factory waste) the pH increased by 2.65 over a distance of approximately 2 mm around the root (scale parameter 0.51 in sigmoid growth). In contrast, the increase of soil pH around roots grown in soil made of pellet FEP particles (virgin material prior to melt processing) was limited to 0.87 and this was observed at greater distance than in soil made of chunk FEP particles (scale factor 0.65, Figure 5E). Application of the Likelihood Ratio tests (χ^2^=1612.45, p < 0.001) showed this trend was highly significant. Chunk FEP particles also produced a more heterogeneous chemical environment with variance increasing with distance from the root (Figure 5F) and therefore equated more closely to the properties of natural soils.

**Figure 5.**
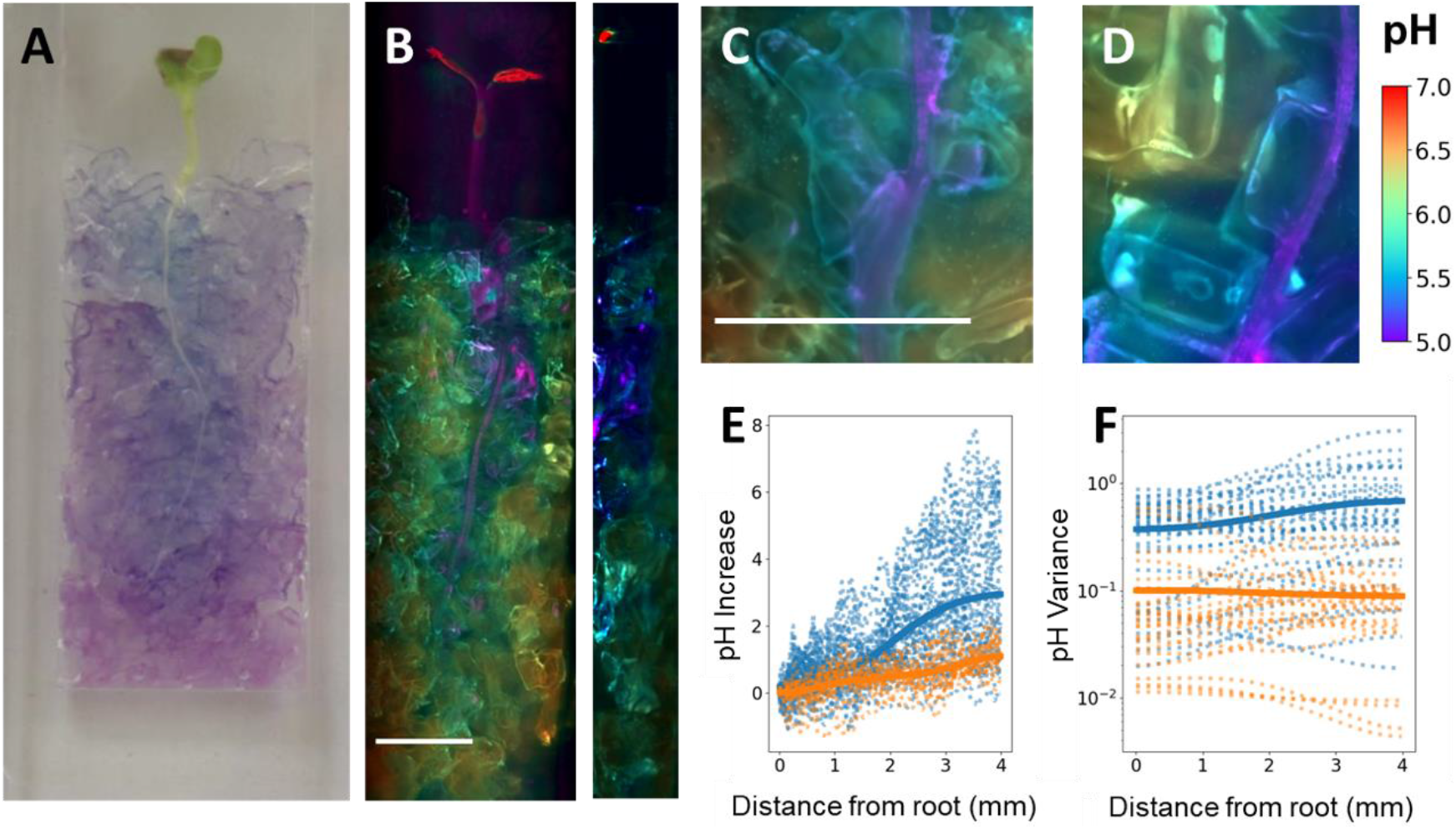
Smart particles reveal how interaction between root exudation and particle type affects the chemical heterogeneity of soils. (A) Seedlings grown in smart soil induced changes in colours of particles due to acidification. (B) Using our multispectral light sheet microscope, pH values were estimated from each optical sections to reconstruct a 3D maps of pH distribution in the inner soil (max projection on the left, cross section on the right, scale bar of 4 mm). Data was acquired on two types of soils, chunk FEP particles (C) and pellet FEP particles (D) represented with a scale bar of 1 mm. (E) Plants grown in chunk FEP particles (blue) induced a more dramatic pH increase with distance from the root than those observed when plants grew in pellet FEP particles (orange). (F) In the soil made of chunk FEP particles, the variance of the pH signal increased with distance from the root, whereas in the case of pellet FEP particles, the variance was reduced. Dotted lines indicate raw data extracted from the image and plain lines indicate the fit of the logistic function. Non-linear mixed effect models showed the pH response observed in the two soils was statistically different (χ^2^=1612.45, p < 0.001).

## Discussion

Soils segregate solids, liquids and gases into a myriad of micro-habitats with distinct physical and chemical composition. Soil organisms have evolved over millions of years to exploit this heterogeneous resource^29^ adapting through specialisation of cell functions and the emergence of complex ecological functions^30,31^ to compete for niche space. Developing artificial media that match or exceed soils’ ability to sustain biological diversity is a scientific challenge that cannot be met using traditional cultivation techniques based on hydrogels, liquids or aerosols.

This study investigated the use of new functional materials to not only mimic complex physical properties of soils, but also enable monitoring of chemical properties that aid understanding and, downstream, potentially allowing control of biological activity within the substrate pores. We proposed a three-steps approach for fast development of soil prototypes using factory waste. Synthesis of functional polymers is first optimised without the core for characterisation of chemical, physical and optical properties. Synthesis is followed by fast particle prototyping using Fluorine-Fluorine interaction^22,23^ to coat FEP particles and allowing the blending of polymers or the recovery of the polymers tested for reuse. Production of the final prototype is then achieved by graft polymerisation for long term stability of the particle^32^. We identified key chemical (surface attachment and stability, ion exchange, functionalisation), optical (refractive index, fluorescence imaging and light transmittance), mechanical (particle size distribution, water retention), technological (reusability, low cost) and environmental (use of un-recycled waste^33^) requirements for the development of such new materials.

Roots are known to release various organic acids to drive soil pH down and solubilise mineral ions adsorbed on soil particles^34^. Results obtained with the smart soil showed acidification occurred within a few millimetres from the root surface which confirms earlier studies^35^, but utilisation of our new technology allows the characterisation of pH changes induced by root exudation at far higher resolution than previously possible with techniques such as planar optodes or zymography assays, where only data from the soil surface can be obtained^36^. Current non-destructive imaging techniques have not, previous to this study, delivered live observations of chemical or biological activity *in situ* within soil, and promising applications of microfluidics remain unsuitable for the study of complex soil environments^37^.The smart soils revealed that chemical changes induced by the roots are intimately related to the structure of the granular media itself. Surprisingly, finer soil texture introduced more variable soil conditions around the root, perhaps due to reduced diffusion coefficients interacting with variation in exudation patterns.

Although smart soils did not enhance plant growth compared to real soil, their successful utilisation here indicate potential for broader application in science or industry. Material sciences in particular offer huge opportunities to further enhance the capabilities of smart soil particles. Control of water and nutrient retention by smart soil could be obtained using techniques improving the wettability of polymers, for example surface texturing and polymer brush structures ^38,39^. Optimisation of porosity of functional polymers, particle shape and size may be exploited to increase the fraction of small pores and therefore the diversity of habitats available to microorganisms^40^. The roughness of particles has the potential to limit friction and resistance to root growth and penetration^41^ and in the more distant future, soil functions themselves could be diversified. For example, feedback between particles and organisms could be engineered to release growth promoters and nutrients or to allow maintenance of complex microbial communities.

Smart soil technologies are bringing unique new capabilities to monitor and quantify biological and chemical changes of the soil environment induced by plants, and in the future, they could help understand soil biodiversity and promote new forms of indoor farming.

## Materials and methods

### Synthesis and characterisation

Reagents consisted of 2,2,2-trifluoroethyl α-fluoroacrylate (FATRIFE) and hexafluoroisopropyl α-fluoroacrylate (FAHFIP) purchased from Scientific Industrial Application P and M (Russia). Poly(ethylene glycol) methyl ether methacrylate (*M_n_* = 300 g/mol), Nile Blue A (>75%), acryloyl chloride (SOCl_2_, 97 %), methyl ethyl ketone (MEK), acetonitrile purchased from Sigma Aldrich, *Tert*-butyl peroxypivalate (TBPPi, 75%) purchased from Akzo Nobel (Chalons-sur-Marne, France). Nile Blue methacrylate (NBMA) was synthesized according previous reports^25^. Terpolymers were characterized by ^1^H and ^19^F NMR spectroscopies (Bruker AC 400 spectrometer). Size exclusion chromatography (SEC) used 0.1 M LiBr/DMF as the eluent, run with a Varian Prostar (model 210) with PPM standards. Fourier transform infrared spectroscopy (FTIR) analyses were performed in the ATR mode using a Perkin-Elmer Spectrum 1000. The synthesis of poly(FATRIFE-ter-PEGMA-ter-NBMA) terpolymers was achieved by conventional radical polymerization (Figure 2A). A 50 ml round bottom flask with a magnetic stirring bar was charged with FATRIFE (4.94 g, 28.70 mmol), PEGMA (8.60 g, 28.70 mmol), NBMA (29 mg, 0.057 mmol), TBPPi (0.101 g, 0.57 mmol) and 72 ml DMF. The mixed solution was deoxygenated with nitrogen bubbling for 15 mins and then placed in an oil bath at 56 °C. The terpolymerization was stopped after 4 hours by immersing the flask in liquid nitrogen and exposing to air. The crude product was purified by precipitation in water, collected by filtration and lyophilised (81 % yield). Dynamic contact angle measurements were performed with a contact angle goniometer (OCA contact angle system, Neurtek Instruments, Spain), equipped with a video camera and image analyser, at room temperature with the sessile drop technique (6 μl, n=5). Spectral properties of the polymers were characterised using a standard multiplate reader (Varioskan™ LUX Multimode, Thermo Scientific, UK).

### Preparation of FEP particles

FEP particles were obtained from Holscott (Grantham, UK) in the form of pellets (virgin material used for extrusion) and chunks (factory waste). Chunk FEP particles were sieved so that the size of particles used were smaller than 1.25 mm and larger than 0.25 mm. Both types of particles were treated in oxygen plasma at 100 W for 1 min (HPT-100, Henniker, UK) to remove dust and residues. 0.4 g of poly(FATRIFE-*ter*-PEGMA-*ter*-NBMA) terpolymer was dissolved in 25ml methyl-ethyl ketone (25642.325, VWR, UK) in a 500 ml rotary evaporator flask containing 20 g of FEP at 40 °C to coat FEP particles. The rotation speed was increased progressively during the evaporation of the solvent to reach a speed of 240 rpm. Particles were then dried overnight at 40 °C. To test for coating stability particles were placed in a test tube immersed in water and spun horizontally at room temperature for several weeks. Then, the pellets were filtered and dried in oven for 12 hrs at 40 °C and characterised by FTIR spectroscopy (Spectrum 1000, Perkin-Elmer, France).

### Plant samples

For the experiments Lettuce sativa seedlings (all year round, Sutton seeds, UK) were used. After a surface sterilisation protocol ^15^ the seeds were incubated on the surface of agar gel (water agar 1%) in a Petri dish under 20°C for 24h wrapped into an aluminium foil to block the light. After the germination the seedlings were inserted into the growth chambers on the surface of the soils (smart, agar, sand and bulk). The growth chambers were kept in an incubator under 20°C with 16h bright period (60μmol/mm^2^s) and 8h dark period. The pH change in the smart soil was observable after 4 days respectively. The microscope images were taken after 1 week and the soil comparison measurements were running for 2 weeks to see the biocompatibility of smart soil.

### Quantitative imaging

Quantitative imaging was performed using a Light Sheet Fluorescence Microscope (LSFM)^28^ equipped with a four wavelength-laser source (Vortran Versalase, Laser 2000 Ltd, UK) with wavelength *λ_i_*, *i* ∈ {488 nm, 514 nm, 561 nm and 633 nm} with the imaging arm was fitted with long pass filters of 530 nm, 570 nm, 645 nm and 665 nm, respectively (Thorlabs FGL530, FGL570, FGL645, FGL665). The signal corresponding to each fluorescence signal was recorded in separate images *I_i_*(*x*, *y*), and a measure of the pH was determined based on the pixel intensity of each of the images.

A dataset was assembled to calibrate pH prediction. Image data was collected from 3 experimental runs at 11 different pH levels achieved using MES acid monohydrate and Na buffers at respective pH of 4.97, 5.2, 5.37, 5.58, 5.78, 6.03, 6.18, 6.38, 6.63, 6.8 and 6.97 (Merck 1.06126 and 1.06197). For each pH, 12 images were collected covering all 4 wavelengths. Points of interests were automatically extracted from images obtained at 488 nm excitation and 530 nm emission (Figure S1). These points were then used to collect a dataset describing the relation between pixel intensities at all wavelengths and pH values. Regressions were established between pH and image intensity ratios 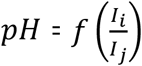 (Figure S2). To further improve predictions, neural network models of the type *pH* = *f*(*I_i_*, *x*, *y*), were developed using Multi-layer Perceptron regressor with rectified linear unit function and trained using a Limited-memory BFGS optimiser. The final neural network model contained 3 layers of 12 neurons each (Figures S3 and S4).

### Computation and statistical analysis

Computations were performed using the Scikit-learn library ^42^. Datasets of pixel intensity at known pH was compiled with macros and scripts developed in ImageJ. Statistical analysis was performed using R with analysis of pH variations surrounding plant roots done using the nlme library^43,44^.

## Acknowledgements

The authors thank Akzo and Tosoh Finechemicals Corp. for supplying with *tert*-butyl peroxypivalate and 2-trifluoromethacrylic acid, respectively.

European Research Council (ERC) under the European Union’s Horizon 2020 research and innovation programme (Grant agreement No. 647857-SENSOILS).

Researchers at the James Hutton Institute also receive financial support from the Rural & Environment Science & Analytical Services Division of the Scottish Government.

## Author contributions

Conceptualization: LXD, BA, VL, MPM

Methodology: LXD, BA, VL, MPM, TSG

Investigation: DP, QY, PF, YH, BB, BVT

Funding acquisition: LXD

Project administration: LXD

Supervision: LXD, BA, TSG, TJD, VL, MPM

Writing – original draft: DP, QY, PF, YH, LXD

Writing – review & editing: BA, VL, MPM, TSG, TJD

## Declaration of interests

There is NO Competing Interest.

## Notes

### Competing Interest Statement

The authors have declared no competing interest.

